# Basolateral amygdala dopamine signals behavioural salience across exploration and learning

**DOI:** 10.64898/2026.09.03.749083

**Authors:** Jessica Capece Marsico, Malika Askarova, Olga Sharma, Sangeetha Nandakumar, Catarina Pacheco, Natalia Favila, Sandra Blaess, Sabine Krabbe

## Abstract

Animals must balance exploration with threat avoidance, yet how neuromodulatory signals shape information-seeking states remains unclear. Using fibre photometry recordings in the basolateral amygdala in freely moving mice, we show that dopamine release is selectively elevated during self-initiated exploratory actions, adapts with repeated exposure, and tracks the behavioural relevance of predictive cues in associative learning and extinction. Our results advance current models of amygdala dopamine function by suggesting that it conveys a state-dependent salience signal that highlights behaviourally relevant moments arising from both environmental cues and self-generated exploration, thereby priming amygdala circuits for plasticity.

## MAIN

Animals continually evaluate their surroundings to balance the competing demands of seeking rewards and resources while avoiding potential threats. To adapt and make accurate predictions about the future, they must learn not only from explicit reinforcement, but also actively explore their surroundings to gather information that is independent of current need states. Exploration serves not merely to guide immediate actions, but to reduce uncertainty and build internal models of the environment that support future decision-making^1,2^. Without such information-seeking, animals risk relying on outdated predictions and failing to adapt to changing contexts.

Beyond its canonical role in associative learning^3^, the basolateral amygdala (BLA) encodes shifts between exploratory and defensive states^4,5^, suggesting that it integrates emotional significance with ongoing behaviour. These dynamics imply that BLA activity must be shaped by external signals that reflect the behavioural significance of ongoing events. The neuromodulator dopamine is a prime candidate, as dopaminergic afferents to the BLA promote both reward- and threat-related learning^6–8^ and have recently been proposed to signal behavioural salience related to associative learning^7,9^. Yet it remains unclear how BLA dopamine is engaged during naturalistic exploration, when animals gather information and evaluate potential significance before outcomes occur.

To address this gap, we used fibre photometry in mice to record BLA dopamine release during multiple self-paced exploratory behaviours and associative learning (Extended Data Fig. 1a) with the genetically-encoded fluorescent dopamine sensor dLight1.1^10^ (Extended Data Fig. 1b,c). To examine how BLA dopamine is engaged during self-paced exploratory decisions, we first monitored dopamine release in the elevated zero maze (EZM), a paradigm that places animals in conflict between the drive to explore and the avoidance of exposed spaces (Fig. 1a-b, Extended Data Fig. 2a-b). Mice freely alternated between open and closed arms, spending less time in the exposed regions but moving faster when they ventured into them (Extended Data Fig. 2c-e). Spatial maps of z-scored dLight signal showed localised increase of dopamine release in the open arms, particularly near transition zones to the closed arms (Fig. 1c). Analysis across all mice revealed that transitions into the open arms were consistently accompanied by increases in BLA dopamine, compared to a significant decrease when transitioning into the closed arms, even when speed was matched (Fig. 1d-f, Extended Data Fig. 2e). Dopamine levels also increased during head dips at the edge of the maze, a hallmark of exploratory risk assessment (Fig. 1c-d, g). No corresponding signal dynamics were observed around open or closed arm entries or head dips in GFP-expressing control mice (Extended Data Fig. 1d-e), supporting the specificity of these behaviour-linked dopamine signals (Extended Data Fig. 2g-i).

**Figure 1:**
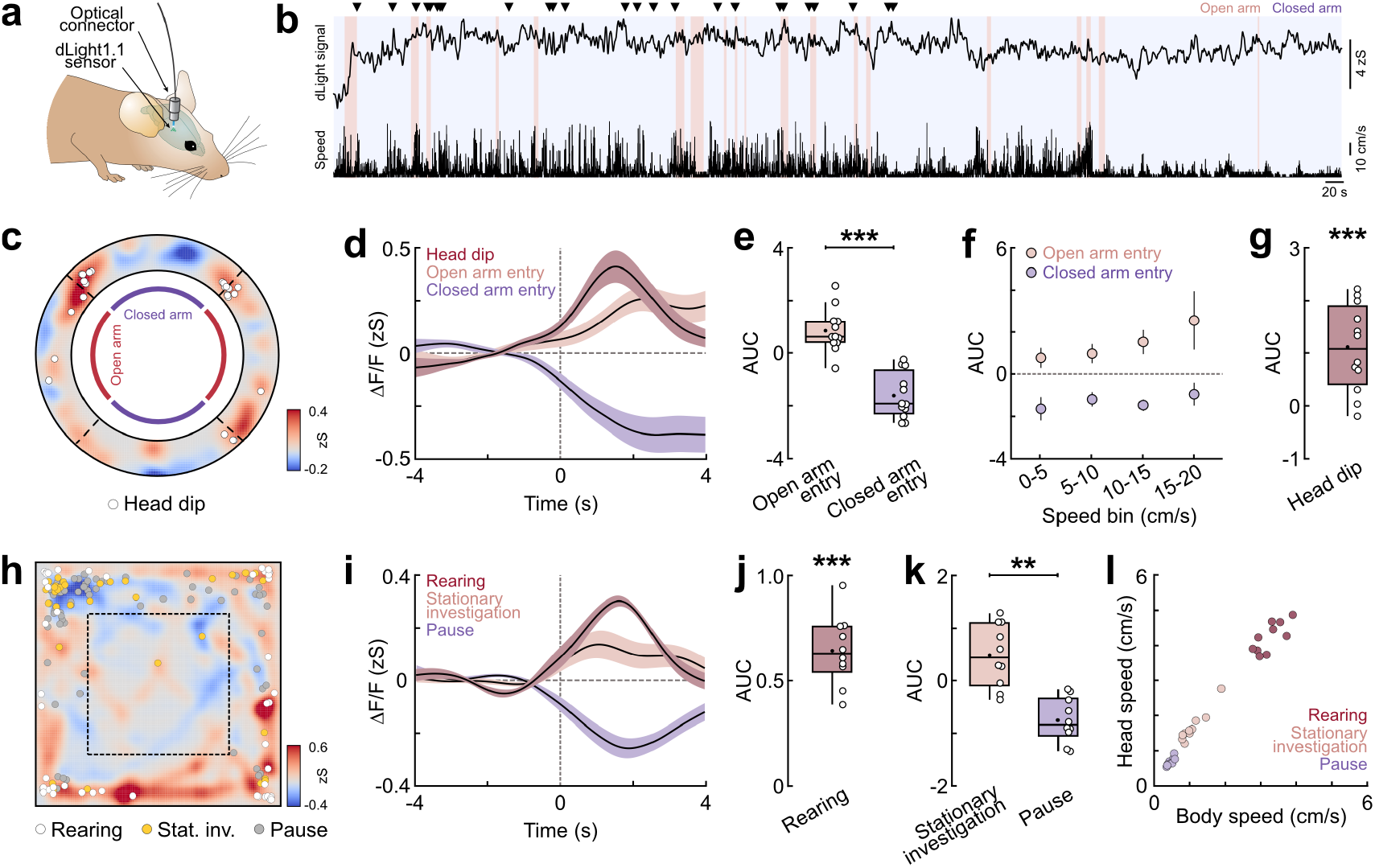
Basolateral amygdala dopamine release tracks exploratory states and transitions. **a** Schematic of the approach used for fibre photometry recordings of dopamine release in the basolateral amygdala. **b** Top, representative example trace of BLA dopamine release in the elevated zero maze (EZM), bottom, corresponding speed trace of the animal, for the entire 20-min session. Colours indicate periods spent in the closed (purple) and open (red) arms (referring to the centre of mass of the mouse), triangles indicate timepoints of head dips. **c** Spatial heatmap of dopamine release in the EZM for the same example mouse. Circles indicate body location (centre of mass) at the timepoint of head dip. **d** Dopamine release time-locked to head dips, open and closed arm entries in the EZM (N=12 mice). **e** Average area under the curve (AUC) for open vs. closed arm entries. Paired t test, *t*(11)=5.716, *p=*0.0001; N=12. **f** Dopamine release across locomotor-speed bins during open- and closed-arm entries. Differences between open and closed arm entries did not depend on locomotor speed, as indicated by the absence of a speed bin × arm interaction. Linear mixed-effects model, Type III ANOVA with Satterthwaite’s approximation; speed bin: *F*(3,71)=1.330, *p*=0.2716; arm: *F*(1,71)=54.02, *p*<0.0001; interaction speed bin × arm: *F*(3,71)=0.651, *p*=0.5847; N=12. **g** Average AUC for head dip events. One-sample t test, *t*(11)=4.651, *p*=0.0007; N=12. **h** Representative spatial heatmap of dopamine release in the open field (OF). Circles indicate body location (centre of mass) at the timepoints of rearing (white), stationary investigation (Stat. Inv., yellow) and pauses (grey). Stationary investigation refers to local exploratory actions, such as floor, wall or corner sniffing, nose-up sniffing, or brief digging-like movements close to the wall. In contrast, pauses reflect locomotor arrest. **i** Dopamine release time-locked to rearing, stationary investigation and pauses in the OF (N=10). **j** Average AUC for rearing events. One-sample t test, *t*(9)=12.40, *p*<0.0001; N=10. **k** Average AUC for stationary investigation vs. pauses. Paired t test, *t*(9)=4.486, *p=*0.0015; N=10. **l** Average body and head movement during pauses, stationary investigation and rearing (N=10). Average traces in d and i are weighted means across animals with s.e.m.; data in f represents mean with s.e.m.; Tukey box-and-whisker plots in e, g, j and k show median values, 25^th^ and 75^th^ percentiles, and min to max whiskers with exception of outliers, dots indicate the mean, circles represent individual animals. \*\**p*<0.01, \*\*\**p*<0.001. Additional details of statistical analyses are provided in Extended Data Table 1.

To further characterise BLA dopamine dynamics during unstructured self-paced behaviour, we recorded release during open field (OF) exploration (Fig. 1h, Extended Data Fig. 3a). Mice preferentially occupied peripheral zones but exhibited variability in overall locomotor activity across animals (Extended Data Fig. 3b-c). BLA dopamine increased robustly during rearing, a highly exploratory behaviour, indicating strong recruitment during active environmental sampling (Fig. 1i-j, Extended Data Fig. 3d). To determine whether increased BLA dopamine during rearing reflected exploratory engagement rather than movement itself, we compared distinct behavioural states occurring during similarly low translational movement. These periods were classified as pauses, reflecting genuine locomotor arrest (translational body movement less than 0.6 cm/s, Fig. 1l), or as stationary investigation when mice continued to display local exploratory actions, including floor, wall or corner sniffing, nose-up sniffing, or brief digging-like movements close to the wall (translational body movement less than 2 cm/s, Fig. 1l). Dopamine increased during these stationary investigatory bouts but decreased markedly during full locomotor arrest (Fig. 1i, k-l, Extended Data Fig. 3e). Thus, BLA dopamine tracked exploratory engagement even in the absence of active locomotion, resulting in different dopamine dynamics across these two distinct behavioural states. Consistent with this dissociation from locomotor activity, locomotor speed explained on average only 1.2% of the within-mouse variance in BLA dopamine signals across the full speed range, decreasing to 0.4% when low-speed periods were excluded (<3 cm/s, removing pauses and stationary investigation to restrict the analysis to active locomotion and minimising the contribution of low-translational movement; Fig. 1l, Extended Data Fig. 3f-i). Furthermore, corner entries and exits, previously linked to transitions between exploratory and non-exploratory states in the OF^4^, were accompanied by modest but non-significant changes in the corresponding direction (Extended Data Fig. 3j-l). Together, these results indicate that BLA dopamine is preferentially recruited during self-initiated exploratory engagement rather than simply scaling with locomotor activity.

To examine the role of BLA dopamine in stimulus-directed exploration, we performed additional release recordings during exploration of an object and a conspecific in an enclosure on the following two days (Extended Data Fig. 4a-d). Because stimulus interactions are inherently directional, we developed an analysis framework to assess dopamine release as a function of the animal’s orientation and proximity to the target (Methods, Fig. 2a-c, Extended Data Fig. 4e). Transforming behavioural data into a target-centred coordinate space revealed that non-selective maps were spatially diffuse (Fig. 2b-c). When frames were restricted to periods in which the mouse was oriented towards the target, dopamine signals became spatially focussed and target-centred (Fig. 2b-c). This orientation filter was used to isolate episodes of target-directed exploration across both paradigms. Within this orientation-defined state, dopamine release scaled with distance from the target. Radial profiles revealed a clear gradient centred on the target for both object and social exploration, with higher dopamine levels in proximal relative to distal zones for both conditions (Fig. 2d-g).

**Figure 2:**
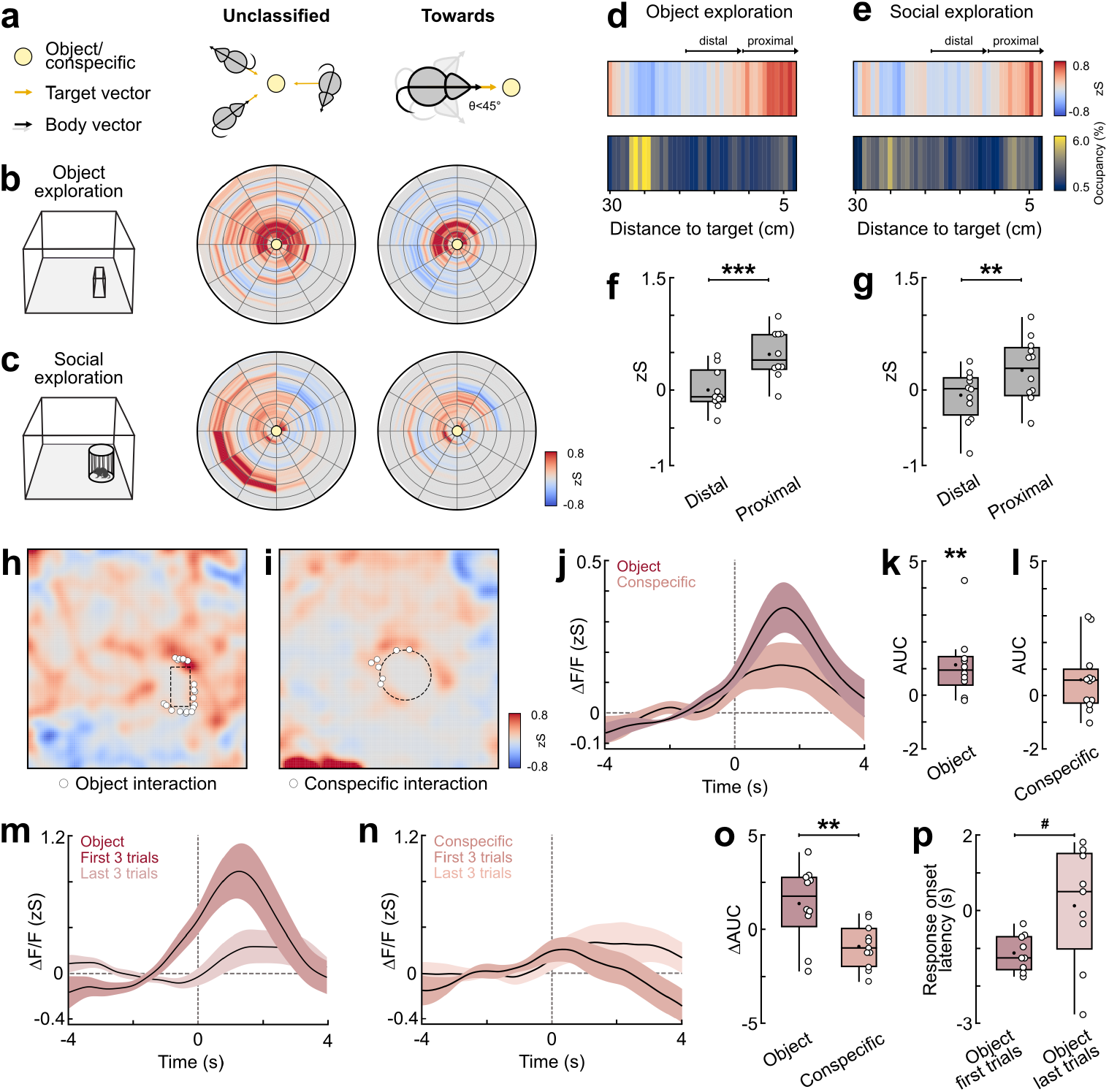
Basolateral amygdala dopamine release scales with proximity during stimulus-directed exploration. **a** Schematic illustration of the vector-based orientation classification method. An orientation angle θ was computed as the angle between the animal’s body vector (yellow arrow) and the target vector (black vector) pointing to the centre of the object or the enclosure (represented as yellow circle). For the “unclassified” category, all behavioural frames were included, regardless of the animal’s orientation. For “towards”, only frames with an orientation angle θ<45° are included. **b** Example polar heatmaps for object exploration for all frames (“unclassified”, left) and selective for “towards” frames (right). Maps are constructed by converting each frame into polar coordinates relative to the stimulus and binning the z-scored signal by angular sector and radial distance from the object’s centre (radius 30 cm). **c** Corresponding example polar heatmaps for social exploration, same mouse. **d** Average one-dimensional maps showing occupancy-normalised dopamine release (top) and spatial occupancy (bottom) as a function of radial distance from the object’s centre (N=10 mice). **e** Same for social exploration (N=12). **f** Average z-score of dopamine release comparing distal (11-19 cm) and proximal (3-11 cm) approach zones for object exploration illustrating the distance-dependent profile of dopamine release. Paired t test, *t*(9)=6.051, *p=*0.0002; N=10. **g** Average z-score of dopamine release comparing distal and proximal approach zones for social exploration. Paired t test, *t*(11)=3.149, *p=*0.0093; N=12. **h** Representative spatial heatmap of dopamine release during object exploration (same mouse as in b-c). Circles indicate location of nose keypoint at the timepoint of close contact with the object. **i** Representative spatial heatmap of dopamine release during social exploration (same mouse as in b-c). Circles indicate nose-to-nose contact with the conspecific through the rods of the enclosure. **j** Dopamine release time-locked to object or conspecific interaction (N=10 for object, N=12 for conspecific). Timepoint 0 defines nose-to-object or nose-to-nose contact, respectively. **k** Average area under the curve (AUC) for nose-to-object events. One-sample Wilcoxon test, *W*=49.00, *p*=0.0098; N=10. **l** Average AUC for nose-to-nose events. One-sample t test, *t*(11)=1.692, *p*=0.1188; N=12. **m** Dopamine release time-locked to the first three and last three nose-to-object contacts in object exploration (N=10). **n** Same for social exploration and nose-to-nose contacts with the conspecific in an enclosure with rods. Only mice with at least 6 trials were included (N=11). **o** Difference in AUC between the first three vs last three trials for object and conspecific contacts. Unpaired t test, *t*(19)=3.144, *p=*0.0053; N=10, object; N=11, conspecific. **p** Response onset for object contact, defined as the first time point at which the signal reached 10% of its peak amplitude (t_10_), relative to contact onset (0 s). Negative values indicate pre-contact onset during close approach. Onset latency was calculated only for mice exhibiting a positive response peak within the predefined response window (0-4 s) for both timepoints (N=9). Paired t test, *t*(8)=2.033, ^#^*p*=0.0765. Average traces in j, m and n are weighted means across animals with s.e.m.; Tukey box-and-whisker plots in f-g, k-l, and o-p show median values, 25^th^ and 75^th^ percentiles, and min to max whiskers with exception of outliers, dots indicate the mean, circles represent individual animals. \*\**p*<0.01, \*\*\**p*<0.001. Additional details of statistical analyses are provided in Extended Data Table 1.

We next examined dopamine dynamics at the point of close interaction (Fig. 2h-j, Extended Data Fig. 4f-i). Dopamine signals increased with the onset of object interaction but showed a more modest modulation during nose-to-nose conspecific contact (Fig. 2j). Quantification revealed a robust enhancement of dopamine during close-contact object exploration relative to baseline (Fig. 2k), whereas conspecific interactions did not differ significantly (Fig. 2l), suggesting that BLA dopamine is more strongly recruited during close engagement with non-social stimuli. We next asked whether these signals remained stable over repeated encounters. Compared to social contacts, dopamine responses to object interactions declined across time (Fig. 2m-o). Object-evoked responses also showed a tendency to become more tightly locked to contact across repeated encounters, whereas response timing remained stable during conspecific interactions (Fig. 2p, Extended Data Fig. 4j). The attenuation of object-evoked dopamine responses, together with a tendency towards increasingly contact-locked response timing, is consistent with a progressive reduction in stimulus novelty or informational value, in line with dopamine signalling of novelty in other brain regions, including the neighbouring caudal striatum^11,12^. Together, these results indicate that BLA dopamine is organised around target-directed exploration and increases with proximity, but is most strongly recruited during close engagement with non-social stimuli and adapts across repeated object encounters.

Having established that BLA dopamine is released upon exploratory engagement and proximity to behaviourally salient stimuli, we next asked whether similar principles apply when animals learn about cues predicting salient events. Therefore, we addressed how BLA dopamine is engaged during associative learning and subsequent extinction (Methods, Extended Data Fig. 1a). During auditory fear conditioning, consisting of five pairings of an auditory cue (CS+, conditioned stimulus) with a mildly aversive foot shock (US, unconditioned stimulus), dopamine release was strongly increased at the time of the aversive US (Fig. 3a-b), with declining amplitudes across repeated pairings with the CS+ (Fig. 3c-d). A significant upregulation during learning could be detected for the CS+ (Fig. 3a-b, e-f), whereas responses to an unpaired CS−control tone remained modest (Fig. 3a-b, g-h). No corresponding cue- or shock-locked signal dynamics were observed in GFP-expressing control mice (Extended Data Fig. 5l).

**Figure 3:**
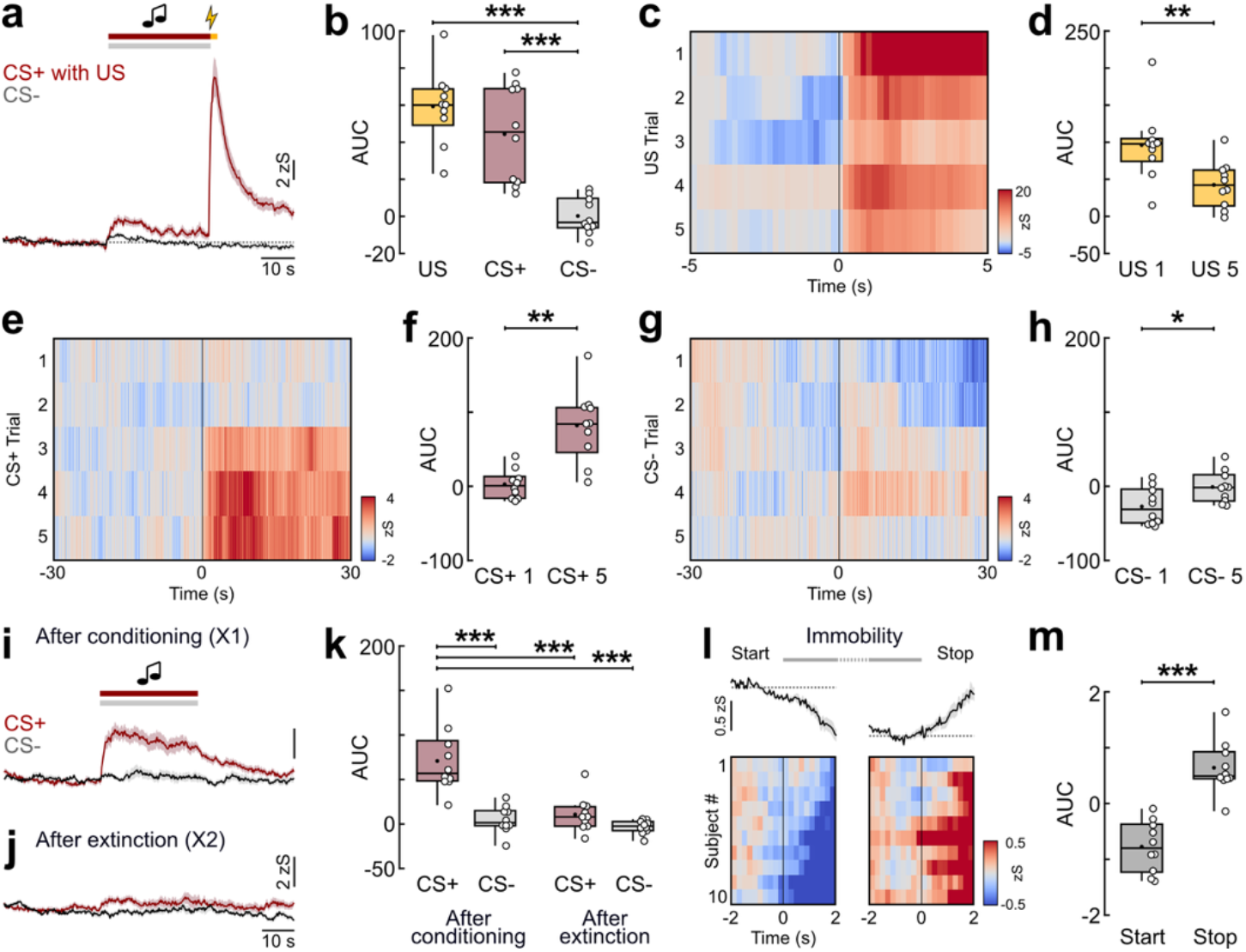
Learned cue relevance shapes amygdala dopamine release during fear conditioning and extinction. **a** Basolateral amygdala dopamine release during fear conditioning for CS+ and US presentations (red) as well as for CS−control cues (black). Shown is the average trace across five trials (N=10 mice). **b** Average area under the curve (AUC) for US, CS+ and CS−. Repeated-measures one-way ANOVA, *F*(2,18)=31.15, *p<*0.0001, followed by Holm-Šídák multiple comparisons: US vs. CS−, *p<*0.0001; CS+ vs. CS−, *p<*0.0001; US vs. CS+, *p*=0.0722; N=10. **c** Heatmap of dopamine release during the aversive US illustrating decline in US evoked signals across five trials during associative learning (N=10). Timepoint 0 indicates US onset. **d** Comparison of US evoked dopamine release during the first and last US presentations. Paired t test, *t*(9)=3.471, *p=*0.0070; N=10. **e** Heatmap of dopamine release during the predictive CS+ demonstrating increase in signals across five trials during associative learning (N=10). **f** Comparison of CS+ evoked dopamine release during the first and last CS+ presentations. Paired t test, *t*(9)=4.151, *p=*0.0025; N=10. **g** Heatmap of dopamine release during the control CS−indicating modest increase in signals across five trials during conditioning (N=10). **h** Comparison of CS−evoked dopamine release during the first and last CS−presentations. Paired t test, *t*(9)=2.301, *p=*0.0470; N=10. **i** Average trace for dopamine release during the CS− and CS+ after conditioning (first presentations in Test/X1 session, 24 h after learning; N=10). **j** Average trace for dopamine release during the CS− and CS+ after extinction (end of X2 session; N=10). **k** Comparison of CS+ and CS−evoked dopamine release over the course of fear expression (“after conditioning”) and after extinction. Repeated-measures one-way ANOVA, *F*(3,27)=19.08, *p<*0.0001, followed by Holm-Šídák multiple comparisons: CS+ after conditioning vs. CS−after conditioning, *p<*0.0001; CS+ after conditioning vs. CS+ after extinction, *p<*0.0001; CS+ after conditioning vs. CS−after extinction, *p<*0.0001; all other comparisons n.s.; N=10. **l** Average traces and heatmap of dopamine release during immobility bouts, averaged across events outside CS presentations in Test/Extinction 1 and Extinction 2 sessions, aligned to immobility start and movement onset (= immobility stop). Mice are sorted by response amplitude upon immobility start (N=10). **m** Corresponding AUC for immobility start and stop. Paired t test, *t*(9)=5.425, *p=*0.0004; N=10. Average traces in a, i and j are means across animals with s.e.m.; traces in l represent weighted means across animals with s.e.m.; Tukey box-and-whisker plots in b, d, f, h, k and m show median values, 25^th^ and 75^th^ percentiles, and min to max whiskers with exception of outliers, dots indicate the mean, circles represent individual animals. \**p*<0.05, \*\**p*<0.01, \*\*\**p*<0.001. Additional details of statistical analyses are provided in Extended Data Table 1.

Across days, BLA dopamine release during the CS+ was low during initial habituation (Extended Data Fig. 5a-c), increased in a high-fear state after conditioning, and declined again upon extinction training – an effect not visible for the control CS−for which dopamine release remained minimal (Fig. 3i-k). Within extinction sessions, CS+-evoked dopamine release progressively declined across repeated cue presentations on the first day, partially recovered at the beginning of the second day, and declined again with continued extinction training (Extended Data Fig. 5d-f). These dynamics indicate that BLA dopamine tracks the behavioural relevance of the CS+ as it acquires and subsequently loses predictive value. Of note, there was no detectable response upon US omission after the CS+ in early extinction training (Fig. 3i), which has been previously observed for dopamine release in nucleus accumbens during aversive conditioning^13^. The temporal profile of BLA dopamine responses across learning days is thus overall consistent with an expectation-modulated teaching signal as suggested previously^14^, and parallels neuronal activity in the BLA^4,15,16^.

Because freezing is a prominent behavioural state during fear learning and extinction, we further examined BLA dopamine dynamics during immobility outside cue periods to isolate state-related dynamics from cue-evoked responses. Release decreased during episodes of immobility and increased when movement resumed (Fig. 3l-m). These dynamics were reliably detectable across habituation as well as early and late extinction sessions individually (Extended Data Fig. 5g-k), and matched previous results of dips in dopamine release during locomotor pauses in the open field (Fig. 1i, k).

Across exploratory and conditioning paradigms, our findings suggest that BLA dopamine marks behaviourally salient epochs with heightened potential for updating and learning. Dopamine release increased during informative external events and self-initiated exploratory behaviours, but was suppressed during exploratory disengagement and declined as predictive information lost behavioural relevance, consistent with dopamine transiently placing BLA circuits in a learning-permissive state. During fear learning, the strong CS+-evoked response after conditioning may therefore reflect an expectation-modulated teaching signal that promotes further learning while predictive information remains behaviourally relevant, whereas its progressive decline during extinction may reflect a reduced need for updating. The absence of a detectable response to US omission further distinguishes this signal from a classical signed prediction error.

Ventral tegmental area (VTA) dopaminergic projections to the BLA are rapidly activated during rewards and punishments and respond to predictive cues in a state-dependent manner^6–8,17^. Recent work shows that BLA dopamine supports identity-specific cue–reward memories without conferring incentive value or acting as a reinforcer^7^, and encodes perceived emotional salience while priming, rather than tracking learning^9^. These findings support a state-dependent salience signal that promotes learning beyond canonical reinforcement frameworks. Our data extend this concept to self-initiated exploration: BLA dopamine was recruited during exploratory states and behaviourally meaningful transitions, but was largely uncoupled from locomotor speed (Extended Data Fig. 3f-i), declined with repeated object contacts (Fig. 2m-p), and remained minimal for the CS−despite matched sensory conditions (Fig. 3). Thus, it appears that BLA dopamine reflects behavioural salience linked to learning opportunities, rather than sensory or locomotor properties *per se*. Consistent with such a role of VTA circuitry in active environmental sampling, activation of glutamatergic medial septum inputs to the VTA increases sniffing, rearing and approach to novel objects, but not to conspecifics or food^18^.

An important question is how dopamine affects BLA microcircuits. Large-scale calcium imaging has shown that BLA neurons forming exploratory ensembles are more likely to be activated by the CS+ after fear conditioning^4^, consistent with the idea of ‘predisposition for learning’ of circuit elements engaged by exploration. This overlap may provide a substrate through which exploratory sampling biases later memory formation. Notably, social interaction recruits different BLA neuronal ensembles^5^, consistent with the weaker dopamine release we observed during social compared with spatial/object exploration, and suggesting differential neuromodulatory recruitment across these behavioural repertoires.

At the BLA microcircuit level, dopamine midbrain inputs preferentially target BLA projection neurons containing D1-type dopamine receptors^19^, but also D1- and D2-expressing inhibitory neurons, including interneurons and intercalated cell clusters^20–22^. Nanoscale anatomical data further suggests that dopamine afferents in the amygdala are organised around receptor-defined microdomains, with dopamine boutons preferentially apposed to D1- or D2-receptor puncta rather than acting exclusively through diffuse volume transmission^23^. Functionally, dopamine suppresses feedforward inhibition and alters projection-neuron excitability^21,24,25^, potentially amplifying strong inputs while dampening weaker ones^26^. Such mechanisms could allow behaviourally salient information to be selectively enhanced while suppressing background activity. Our findings are therefore consistent with BLA dopamine acting as a context- and experience-dependent regulator that transiently places BLA microcircuits in a plasticity-permissive state, promoting the incorporation of meaningful exploratory or predictive information while limiting indiscriminate strengthening during behavioural quiescence or predictability.

Although our fibre photometry measurements are correlational, previous ablation and optogenetic studies establish that BLA dopamine can causally influence exploratory and anxiety-related behaviour^27–30^. Such manipulations, however, do not directly test whether endogenous dopamine transients transiently permit plasticity at behaviourally salient moments, as altering dopamine may itself change the behavioural states and sampling that drive learning. Addressing this mechanism will instead require temporally precise manipulation while monitoring BLA circuit dynamics, for example by combining ensemble-level calcium imaging with optogenetic control of dopaminergic axons^16,31^.

In summary, our findings suggest that BLA dopamine provides a state-dependent signal that marks behaviourally relevant moments during information seeking and predictive learning, biasing amygdala circuits toward updating when new information matters most.

## Supplementary information

Extended Data Figures 1-5

Extended Data Table 1

## Data and code availability

Custom code and full datasets will be made available upon final publication and are currently available upon request from SK.

## Author contributions

Conceptualisation: JCM, SK; Methodology: JCM, SK; Investigation: JCM, OS; Analysis: JCM, MA, SN, CP, NF, SK; Visualisation: JCM, MA, SN, SK; Code and data curation: JCM, MA, SN; Funding acquisition: SB, SK; Supervision: SB, SK; Writing – original draft: SK; Writing – review & editing: all authors.

## Acknowledgements

The authors thank all members of the Krabbe lab for helpful discussions and comments. They thank all staff of the DZNE Animal Facility for excellent animal care, breeding management, and technical assistance throughout this study. They further thank the DZNE Light Microscopy Facility for their support with data acquisition and analysis. pAAV-CAG-dLight1.1 was a gift from Lin Tian (Addgene viral prep #111067-AAV5; http://n2t.net/addgene:111067; RRID: Addgene_111067).

## METHODS

### Animals

All animal procedures were conducted in accordance with the European Directive 2010/63/EU and the German Animal Welfare Act, and approved by the Landesamt für Natur, Umwelt und Verbraucherschutz Nordrhein-Westfalen (LANUV NRW, Germany) before experiments started. Experiments were performed with male and female wildtype mice aged 3-6 months at the time of injection. To minimise surplus breeding, we used DAT-Cre wt/wt mice on a C57BL/6J background (original strain: Jax Stock No. 006660, bred with C57BL/6J Stock No. 000664), with exception of two GFP controls (C57BL/6J breeding, IDs 20739 and 20742). Animals were housed on a 12-hour light/dark cycle at constant temperature (22 °C) and humidity (58-60%), with food and water available *ad libitum*. Mice were kept in ventilated cages with bedding, nesting material and enrichment (including igloo and running wheel, Bio-Serv). Starting a week before surgeries, animals were single-housed to prevent later implant damage. Social stimulus mice were single-housed for one week before behavioural experiments.

### Surgical procedures

Mice were anaesthetised using isoflurane (3-5% for induction, 1-2% for maintenance; Vetfluran, Virbac) in oxygen-enriched air (EverFlow OPI, Respironics) and fixed on a stereotactic frame (Model 1900, Kopf Instruments). Mice received carprofen (0.067 mg/ml, Rimadyl, Zoetis) in their drinking water starting 24 hours before surgery and continuing for 72 hours postoperatively. Lidocaine (4 mg/kg, s.c., Lidocain 0.5%, Pharmarissano) and ropivacaine (2 mg/kg, s.c., ROPIvacain, B.Braun) were administered preoperatively for local analgesia under the scalp. To protect the eyes during surgery, an ophthalmic cream (Vitamycin, CP Pharma) was applied and eyes covered with black foil to shield them from light exposure. Rectal temperature was measured and maintained at 36 °C using a feedback-controlled heating pad (FHC), and respiratory rate (1-2 Hz) was monitored throughout the entire surgery. After exposing and levelling the skull, a 0.5 mm diameter craniotomy was drilled over the left amygdala. AAV2/5-CAG-dLight1.1 (ca. 200 nl, 111067-AAV5, Addgene) or AAV2/5-CAG-EGFP (ca. 200 nl, v24-5, VVF Zürich) was unilaterally injected into the BLA using a motorised precision micropositioner (MDS-1, Narishige) and pulled glass pipettes (tip diameter about 20 µm) connected to a Picospritzer III microinjection system (Parker Hannifin Corporation) at the following coordinates from bregma: AP −1.65 mm, ML −3.2 mm and DV 4.5 mm below the cortical surface. To correct for biological variability of skull/brain size, stereotaxic coordinates were scaled according to individual bregma-lambda (BL) distance (AP_corrected_= (BL/4.2) × AP_atlas_+ 0.2, based on the standard BL distance of 4.2 mm according to the 5^th^ edition of *Paxinos and Franklin’s the Mouse Brain in Stereotaxic Coordinates*^32^). Next, an optical fibre was implanted above the virus injection site. Fibres (400 µm diameter, 0.5 NA) were custom-made using ceramic ferrules (CFLC-440, Thorlabs) with 400 µm diameter multimode fibre (FP400URT, Thorlabs) and fibre polishing film (Thorlabs). Output was measured prior to implantation, only fibres with transmission efficiencies in the range of ~75-85% were implanted. After slowly lowering the fibre into the brain, it was fixed to the skull using UV light-curable glue (Loctite 4305, Henkel). The skin was sealed with Scotchbond (3M). Vetbond (3M) and dental cement (Paladur, Kulzer) were used to seal the skull. A custom head bar was embedded into the dental cement to allow for later head-fixed attachment of patch cable for fibre photometry recordings. Fibres were covered with tubing to avoid scratching in the recovery period.

### Behaviour

All behavioural testing was performed during the light phase. Imaging of dopamine release in freely moving animals was performed approximately four weeks after surgery. Prior to experiments, mice were handled for two days (5 min/day), followed by three days of handling with habituation to head fixation and fibre connection to the patch cable (5-10 min/day). To assess dopamine release during self-paced and stimulus-driven exploration, as well as during fear learning and extinction, mice underwent behavioural paradigms in the following order (one per day, see Extended Data Fig. 1a): elevated zero maze (EZM), open field (OF), object exploration (OE), social exploration (SE), followed by aversive conditioning consisting of habituation (Hab), conditioning (FC), and two extinction sessions (X1/X2). Mice were briefly head-fixed on a running wheel before and after each behavioural paradigm to connect and disconnect the optical patch cable. Mice were returned to their home cage immediately after the experiment.

### Exploratory paradigms

Spontaneous and stimulus-driven exploration were tested in a sound-attenuated Isolation Cubicle XL (Ugo Basile, custom design) under consistent conditions: cherry/almond scent (acetophenone, Sigma), light intensity of 23 lux with infrared illumination (Thorlabs, Item No. LIU850A, installed at each corner of the cubicle ceiling), and a fan-induced background noise of 45 dB. Self-paced exploration was monitored for a duration of 20 min on four consecutive days. *Day 1*: Spontaneous exploration was first assessed in the EZM (TSE Systems, E-341012-M-W-IRT), which consisted of two open arms (each with a 0.5 cm ledge) and two closed arms (arm width: 5.5 cm, wall height: 11 cm). The maze had an outer diameter of 46 cm and was elevated 40 cm above the base of the cubicle. The mouse was placed in one of the closed arms and then allowed to explore freely. *Day 2*: The OF test was conducted in a square arena (TSE Systems, E-302050-OF-W-IR-500; 46 × 46 cm, wall height: 40 cm). Each mouse was placed in one corner of the arena and allowed to explore freely. *Day 3*: In the same OF arena, mice were exposed to an object for OE (*Duplo* block, height: 13.4 cm, width: 6.3 cm, depth:3.1 cm) positioned slightly off-centre toward the lower right quadrant. *Day 4*: In the same way, SE was assessed in the OF arena by introducing a stimulus mouse (sex-, age- and strain-matched with the experimental mouse, but no littermate) inside an enclosure (Noldus, MMSOC-W001-V10, diameter: 10 cm, height: 20 cm). The rod design of the enclosure allowed for auditory, visual, olfactory as well as limited tactile interactions. Each arena was thoroughly cleaned with distilled water between experimental animals.

### Aversive conditioning

Associative learning was assessed using a four-day discriminatory auditory fear conditioning paradigm conducted across two distinct contexts. Context A (habituation and extinction) consisted of a clear cylindrical chamber (diameter: 30 cm, height: 35 cm) placed in a dimly-lit environment (~30-50 lux) in the presence of ethanol odour. The chamber was fitted with a smooth white floor insert covered with a bit of each animal’s home cage bedding material. Context B (conditioning) was square-shaped with black rear and side walls and clear front wall (30 × 30 cm, height: 35 cm), placed into bright light conditions (~300 lux) in the presence of acetic acid scent. The chamber floor consisted of a grid for foot shock delivery placed over a white base. All sessions took place in a sound-attenuated Multi Conditioning System box (MCS, TSE Systems) equipped with infrared lighting, fan-induced background noise of 50 dB, and an overhead speaker (MF1-S, Tucker-Davis Technologies) for auditory stimulus presentation. Between mice, contexts were cleaned with distilled water and the respective odour. *Day 1* – Habituation: Mice were habituated in context A for 20 min. Following a 2 min baseline, five presentations of two pure tones each were delivered in alternation: a 6 kHz and a 12 kHz tone (30 s total duration, consisting of 200 ms pips delivered at 1 Hz, 75 dB). Stimuli were presented in alternating order with pseudo-random inter-trial intervals ranging from 30-120 s. *Day 2* – Conditioning: Mice underwent conditioning in context B (20 min). Following a 2 min baseline, 5 CS+ presentations (6 kHz) were each paired with a 2 s 0.65 mA foot shock (unconditioned stimulus, US) delivered by the MCS system immediately after tone offset. Interleaved with the CS+/US presentations were 5 CS−control tones (12 kHz). Stimuli were presented with inter-trial intervals ranging from 30-120 s. *Days 3 and 4* – Extinction: Mice were placed in context A for cue-memory retrieval and extinction learning. After a 2 min baseline, 5 CS−tones were presented, followed by 15 CS+ cues without reinforcement (inter-trial intervals 30-120 s). Day 3 is considered ‘Test/Extinction 1’ or X1, and Day 4 ‘Extinction 2’ or X2. Each extinction session lasted 30 min.

### Data acquisition

Behavioural videos were collected at 40 Hz using a top-mounted, infrared-sensitive camera (AVT Stingray II) with the CinePlex digital video recording system (CinePlex Studio, CPX/V3-S, version 3.12.0.1, Plexon Inc) at 640 × 480 px. The Multi Conditioning System 2.0 software (Open Field/Activity and Fear Conditioning Extended Advanced packages, TSE Systems) was used to trigger start/stop of the recording as well as external stimuli (auditory cues, foot shock) via TTL pulses. A multi-input/output processor (RZ6, Tucker-Davis Technologies) recorded timestamps from all acquisition devices via TTL pulses with the OpenEx software (version 2.32.0, Tucker-Davis Technologies), ensuring precise synchronisation to a shared master clock. Timestamps were collected for session start/stop markers, external cues, as well as all behavioural video and fibre photometry frames. Auditory stimuli were generated using the RZ6 processor with RPvdsEx software (version 96, Tucker-Davis Technologies).

### Fibre photometry

Given the sparse dopaminergic innervation and overlapping noradrenergic inputs that reach the BLA, dLight1.1 was selected for its moderate affinity (~330 nM), large dynamic range (~230% ΔF/F), and ~70-fold selectivity for dopamine over noradrenaline^10,33^. Fluorescence signals were recorded using a custom-built fibre photometry system (INSS Independent NeuroScience Services) configured for dual-wavelength excitation and emission. Excitation light was delivered in an interleaved fashion: 405 nm (isosbestic, dopamine-insensitive control) and 470 nm (dopamine-sensitive excitation of dLight1.1), using LED light sources (M405FP1 and M470F3, Thorlabs) and excitation filters centred at 410/10 nm and 470/10 nm, respectively. Light was collimated and combined via dichroic mirrors, then focused through a 20× Olympus Plan Fluorite objective (0.5 NA, 2.1 mm WD) via a patch cable (Thorlabs, M119L03) connected to the implanted 400 µm core optical fibre targeting the BLA. Fluorescence emitted by dLight1.1 (~525 nm) travelled back through the same fibre and was separated from excitation light using a dual bandpass emission filter centred at 523/25 nm and 610/25 nm, rejecting out-of-band wavelengths. The filtered signal was captured using a Basler acA1920-155um monochrome USB3.0 camera running at 40 Hz (20 ms exposure time), operated via Pylon Viewer 7.1.011271 (Basler AG). LEDs were triggered using the RPvdsEx software and interleaved at 40 Hz, resulting in an effective sampling rate of 20 Hz per channel. LED power was individually adjusted depending on transmission efficiency of the implanted fibre and set to ~150 µW at the implanted fibre tip prior to each experiment.

### Histology

At the end of experiments, mice were transcardially perfused for analysis of fibre placements and AAV expression. To this end, mice were first injected with a lethal mixture of ketamine (250 mg/kg, i.p., Ketamin 10%, Serumwerk Bernburg AG) and medetomidine (2.5 mg/kg, i.p., Dorbene, Zoetis) and then perfused with 4% paraformaldehyde (PFA) in phosphate buffered saline (PBS). To prevent clotting, heparin (50 µl of 5000 I.E./ml, B.Braun) was administered to the left ventricle before perfusion with 4% PFA. Brains were stored in 4% PFA overnight and then transferred to PBS. To confirm correct fibre placement and AAV expression, 100 μm coronal sections of the amygdala were sliced using a vibratome (Leica, VT1000S) and processed for free-floating immunofluorescence. Sections were washed three times in PBS for 10 minutes at room temperature (RT), then blocked in 10% normal goat serum (NGS, Abcam, ab7481) in 0.5% Triton X-100 (Sigma) in PBS (PBST) and incubated for 2 h at RT. Subsequently, slices were incubated with primary antibodies (chicken anti-GFP, 1:1000, Thermo Fisher Scientific, Cat# A10262, Lot# 2407370; rabbit anti-TH, 1:750, Merck Millipore, Cat# 657012, Lot# 3900049) in 1% NGS in 0.5% PBST for 24-48 h at 4 °C. After washing three times for 10 min at RT in 0.1% PBST, secondary antibodies (goat anti-chicken Alexa Fluor 488, 1:750, Thermo Fisher Scientific, Cat# A11039, Lot# 2420700; goat anti-rabbit Alexa Fluor 568, 1:750, Thermo Fisher Scientific, Cat# A11011, Lot# 2782620) were applied in 1% NGS in 0.5% PBST for 12-24 h at 4 °C. Finally, sections were washed twice in PBS for 10 min at RT and then incubated with Hoechst 33258 (10 µg/ml, Thermo Fisher Scientific, Cat# H3569) in PBS for 10 min to stain cell nuclei. After three washes in PBS for 10 min at RT, sections were mounted on glass slides and cover-slipped with Aqua-Poly/Mount (Polysciences). Images were acquired with an AxioScan.Z1 slide scanner (Carl Zeiss AG), equipped with a 10× air objective (Plan-Apochromat 10×/0.45) and excitation laser lines at 353 nm, 493 nm and 577 nm using ZEN blue software (version 3.10, Carl Zeiss AG). Sections were matched against a mouse brain reference atlas^32^ to verify AAV expression and fibre placement within the BLA.

### Behaviour analysis

#### Pose estimation

Pose estimation was performed using DeepLabCut version 2.2.2^34^. All networks used a ResNet-50^35^ backbone and were trained for up to 500,000 iterations, with 95% of frames used for training and 5% for testing. For single-animal tracking, three separate networks grouped by context shape were trained, thereby accommodating habituation/extinction (circular), OF/OE (square), and fear conditioning days (square with grid). A fourth, multi-animal network was trained for SE to distinguish the experimental mouse from the social conspecific. Each network was trained independently using a different number of manually labelled frames, selected across multiple videos. Total number of frames and videos labelled for each network was as follows: *circular network:* 279 frames from 93 videos; *square network*: 186 frames from 62 videos; *square with grid network*: 285 frames from 57 videos; *SE network:* 280 frames from 28 videos. For each frame, 14 anatomical landmarks were manually annotated on the experimental mouse: nose, left and right ear base, neck, upper spine, middle spine, tail base, tail middle, tail tip, left and right forepaws, left and right hind paws, and the base of the implanted optic fibre. In the SE paradigm, 13 corresponding landmarks were also annotated on the stimulus conspecific, excluding the fibre base. After inference, x/y coordinates with likelihood <0.9 were discarded and replaced with the most recent high-confidence position. Coordinates outside predefined spatial boundaries of behavioural contexts (±10% margin) were also excluded. Gaps of up to 10 consecutive frames were linearly interpolated. Coordinates were converted to centimetres using a fixed scaling factor (46 cm = 400 px) and down-sampled from 40 Hz to 20 Hz. Tracking in the EZM was performed using CinePlex Studio software with the CinePlex ‘Tracking and Behavior’ packages (CPT-V3 and CPB-V3, Plexon Inc). Similar pre-processing steps were applied: position data were interpolated (≤10-frame gaps), converted to centimetres, down-sampled to 20 Hz, and filtered to exclude frame-to-frame displacements >10 cm. Speed was calculated as the framewise Euclidean distance multiplied by the sampling rate and thresholded at 50 cm/s to remove high-velocity artefacts.

#### Event detection

Behavioural events were extracted using task-specific pipelines. In the EZM, zone transitions were derived from Plexon tracking software. In the OF, zones were extracted from DeepLabCut centre of mass coordinates using custom Python scripts. A central square was defined per animal with an 80 px inward margin of the arena (leaving a 9.2 cm wall area), and a 40 px buffer (4.6 cm) was applied to its four corners to capture transition zones between corner/centre. Transition segments were labelled based on direction (corner entry, corner exit, or ambiguous) by analysing the preceding and following zone identity and zone bouts labelled as ambiguous and/or shorter than 0.5 s were excluded from the analysis. Head dips were manually annotated, rearing was detected with BehaviorDEPOT v1.6^36^ and results manually curated. Pauses, stationary investigation (both ≥0.5 s) and immobility (≥2 s) were identified based on low displacement (3.5 px/0.4 cm) across five body points (left and right ear base, upper and middle spine, tail base), followed by manual classification: pauses/immobility were defined as full locomotor arrest; in contrast, bouts during which mice continued to display local exploratory actions, including floor, wall, or corner sniffing, nose-up sniffing, or brief digging-like movements close to the wall/corner, were classified as stationary investigation. For nose-to-object contacts in OE, a rectangular zone (50 px/5.75 cm width, 86 px/9.89 cm height) was defined around the object centre. The animal was considered to contact the object when the nose coordinate transitioned from the outside to inside this zone. Nose-to-nose contacts in SE required the experimental animal to move towards the enclosure, with a threshold for transitions defined as round zone around the target (60 px/6.9 cm from the centre). Of these transitions, nose-to-nose contacts were identified manually. Contacts happening within a 2-s window after the previous event were excluded for both paradigms to reflect approach behaviour immediately before contact.

#### Vector-based orientation classification

To quantify how dopamine release varies as a function of orientation and proximity to the target, a frame-by-frame classification based on geometric vector analysis was applied. Each timepoint was segmented into one of three mutually exclusive categories: *sideways, towards*, or *away*, based on the angle between the body axis of the animal and its directional relationship to the target.

*Definition of vectors –* Two vectors were computed for each frame:

1.Body vector representing the facing direction of the animal, defined as:

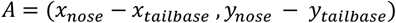

2.Target vector defined as:

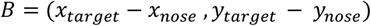

Here, the horizontal (X-axis) is denoted as the component of vector *A* by *a*_*x*_ and the vertical (Y-axis) component by *a*_*y*_. Similarly, B = (*b*_*x*_, *b*_*y*_), where *b*_*x*_ and *b*_*y*_ are its X and Y components.

*Angle computation –* The angle θ (theta) between vectors was calculated using the cosine angle formula:

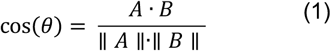

1. The dot product of the two vectors is:

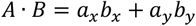
2. Euclidean norms (i.e., length of vectors):

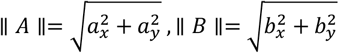

The resulting cosine value was passed through the inverse cosine (arccos) function to recover the angle in radians, which was then converted to degrees using the standard conversion factor:

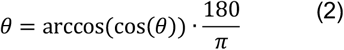

Frames were categorised into three orientation classes based on the angle θ: frames with θ less than 45° were classified as *towards* the target; those with angles between 45° and 135° were labelled as *sideways*; and frames with θ greater than 135° were considered *away* from the target. This classification allows to distinguish between direct engagement, lateral ambiguity, and disengagement from the target. *Towards*-oriented frames were used to isolate target-directed exploration.

*Distance binning and occupancy normalisation –* For each frame, the Euclidean distance between the nose and the target centre was calculated and converted to centimetres. The space around the target was then linearly divided into contiguous distance bins of fixed width (~0.6 cm) spanning 30 cm away from the centre of the target.

For each bin, an occupancy-normalised dopamine signal was calculated:

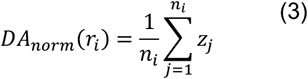

where:

*r*_*i*_is the i-th radial bin (i.e. distance range), *z*_*j*_ is the z-scored ΔF/F value for frame *j*, and *n*_*i*_=is the number of frames in bin *r*_*i*_

For summary comparisons, the bins surrounding the target were divided into two distance-defined regions of equal width. Since distances were calculated from the centre of the target (object or enclosure), the innermost 3 cm fall within the physical boundaries of the target itself, making it physically impossible for the nose to occupy that space. To account for this, the first 3 cm were excluded. The proximal zone was thus defined as the 3-11 cm range, and the distal zone as the 11-19 cm range. This one-dimensional analysis yielded a distance-dependent profile of dopamine release, restricted to frames in which the animal was oriented towards the target.

#### Fibre photometry analysis

Denoising, photobleaching correction, motion correction and normalisation were performed using two different approaches, tailored to the structure of different paradigms.

#### Processing steps exploratory paradigms

In exploration paradigms, behaviour unfolds continuously without predefined trial boundaries. To preserve the temporal integrity of the signal and correct for gradual fluorescence decay, the entire recording was processed using the pMAT v1.3 toolbox^37^. Both the dLight (signal) and isosbestic trace (control) were first denoised using LOWESS smoothing to remove high frequency noise. Motion correction was performed by fitting a global linear regression between the signal and the isosbestic control channel and scaling the signal. The scaled control was calculated as:

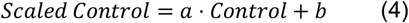

where *a* is the slope, and *b*, is the intercept of the regression line. Then, ΔF/F was computed as:

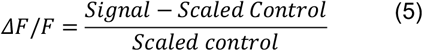

To facilitate cross-subject and cross-condition comparisons, the resulting traces were then normalised across the whole session by computing the z-score as:

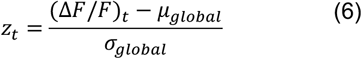

where: µ_global_ is the trace mean of the entire session, and σ_global_ the standard deviation of the trace. Peri-event time heatmaps (PETHs) were generated by aligning z-scored traces to exploratory behavioural events (e.g. head dips). Traces were baselined by subtracting the mean value of a predefined pre-event window from the event trace.

#### Processing steps fear conditioning paradigm

For fear conditioning sessions, given the trial-structure nature of the paradigm, denoising, photobleaching correction, motion correction and normalisation were performed on a per-trial basis using custom Python scripts following PMAT processing steps^37^. Each trial was aligned to a stimulus onset (e.g. tone onset) and included a predefined baseline and event window. After applying LOWESS smoothing to both the signal and isosbestic control traces, a linear regression was used to fit the isosbestic control to the signal in each trial as in equation (4), separately for the event and baseline window. Then, ΔF/F was computed as defined in equation (5), separately for the event and baseline window. Each trial was then zeroed to the first trial point, further correcting for stability and photobleaching. Finally, the traces were standardised using the baseline period:

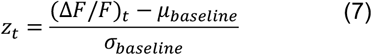

µ_basline_ is the baseline *mean*, and σ_basline_ the baseline *standard deviation*. This trial-wise normalisation accounts not only for photobleaching but also for trial-to-trial variability and is well suited for discrete, cue-locked paradigms^37^.

#### Group-level traces

For event types with variable trial counts across subjects, individual z-scored ΔF/F traces were first aligned to each event and averaged within each animal, producing a single representative trace per mouse. To obtain group-level traces, a weighted mean was computed across animals, where each animal’s contribution was scaled according to the number of valid trials available for that event. This approach ensures that animals with more reliable data have proportionally greater influence on the final average, while still preserving between-subject variability.

The weighted mean was computed as:

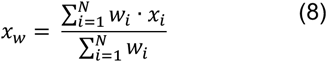

where:

*x*_*i*_ is the mean trace of mouse *i*, and *w*_*i*_ is the number of trials available for mouse *i*.

The weighted variance was then calculated using the standard population formula:

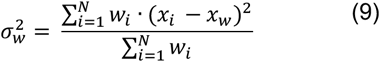

To estimate between-subject variability, the standard error of the mean (SEM) was calculated:

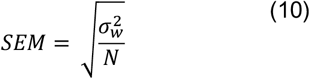

where *N* is the number of animals.

For the fear conditioning paradigm, where all animals had the same number of trials per event type, group-level z-scored ΔF/F traces were computed using a simple arithmetic mean across animals.

#### Event-based analysis

To quantify dopamine release around behavioural events, signals were measured as the area under the curve (AUC) of defined post-event windows compared to the baseline. AUCs were computed in Python (NumPy trapz) and averaged per animal before statistical testing. For exploratory events (head dips, rearing, stationary investigation, pauses), spatial transitions (open vs. closed arm entries, centre vs. periphery), and stimulus-guided exploration bouts (object/conspecific contact), baseline and event AUC windows were set to 4 s. When quantifying signals upon immobility start/stop across fear conditioning days, 2-s windows were used, and only bouts for which the entire peri-event analysis window fell outside CS presentation periods were included. For CS presentations and the aversive US, 30-s and 5-s windows were applied for AUC analysis, respectively.

To quantify changes in object- and conspecific-evoked dopamine responses across repeated interactions, traces were averaged within each mouse across the first three and last three contacts. Response magnitude was quantified as AUC within the predefined 0-4 s post-contact window as described above. To assess response timing, the peak response was identified within the same window for each mouse and condition, and response onset latency was defined as the first time point on the rising phase at which the signal reached 10% of the peak amplitude (t_10_), measured relative to contact onset (0 s). Negative values therefore indicate response onset during the close approach preceding contact. Onset latency was quantified only for mice exhibiting a positive response peak within the predefined response window for both the first and last three approaches.

### Statistical analysis and data presentation

In all figure legends, ‘N’ indicates the number of animals. In total, photometry data from N=13 dLight1.1 (7 male, 6 female) and N=6 GFP mice (3 male, 3 female) with confirmed fibre placement in the BLA was collected. Individual recording days were excluded if major motion artefacts prohibited reliable analysis of fluorescence signal. The resulting ‘N’ number reflecting the number of mice for each behavioural paradigm is stated in each figure legend and in Extended Data Table 1. Sample sizes were chosen based on current standards in the field.

Statistical analyses were carried out using Prism 11 (GraphPad Software) and R (R 4.1.0, RStudio 2023.12.1). All datasets were tested for Gaussian distribution using a Shapiro-Wilk normality test. If the null hypothesis of normal distribution was not rejected, paired or unpaired t-tests, as well as one-sample t-tests against a hypothetical mean of zero, were used as appropriate. A repeated-measures one-way analysis of variance (ANOVA) was used for comparing more than two normally distributed datasets. *Post hoc* multiple comparisons were performed using the Holm-Šídák correction. If the null hypothesis of normal distribution was rejected, pairwise comparisons were performed using a Wilcoxon matched-pairs signed-rank test. One-sample tests were done with a Wilcoxon signed-rank test against a hypothetical median of zero. To determine whether arm entry-evoked dopamine release differed between open and closed arm entries as a function of locomotor speed, AUC values from individual entries were analysed using a linear mixed-effects model. ‘Speed bin’, ‘arm type’ and their interaction were included as fixed effects, and ‘animal’ as a random intercept to account for repeated observations within animals. Fixed effects were evaluated using a Type III ANOVA with Satterthwaite’s approximation. A statistical significance threshold was set at 0.05, and significance levels are presented as \**p*<0.05, \*\**<* 0.01 or \*\*\**p*<0.001 in all figures.

Contrast and brightness of representative histology example images were minimally adjusted equally across the entire image using ImageJ (version 2.1.0/1.53c). Behavioural frames were down-sampled to 20 Hz as explained above. Spatial release heatmaps were normalised by occupancy. Tracking coordinates were grouped into 3 × 3-px spatial bins, and the mean dLight z-score for each occupied bin was calculated by dividing the sum of all z-scores assigned to that bin by the number of contributing samples. Unoccupied bins were assigned a value of zero. The resulting two-dimensional maps were smoothed using a Gaussian filter with a standard deviation (σ) of three spatial bins, truncated at 3σ. For polar release maps, nose positions were transformed into polar coordinates relative to the stimulus centre as described above, and the surrounding space was divided into radial bins of 3 px and angular bins of 30°. For each occupied radial-angular bin, the mean dLight z-score was calculated as described above for spatial release heatmaps, and unoccupied bins were assigned a value of zero. The resulting two-dimensional polar maps were smoothed using a Gaussian filter with a σ of one bin along each dimension, corresponding to 3 px radially and 30° angularly. Averaged dopamine release traces are displayed as mean with s.e.m., using the arithmetic mean across animals for fear conditioning and a weighted mean of LOWESS-smoothed traces weighted by the number of trials contributed by each animal for exploratory behaviours. For figure display, dopamine release heatmaps and traces for CS and US presentations were down-sampled to 5 Hz. Tukey box-and-whisker plots show median values, 25^th^ and 75^th^ percentiles, and min to max whiskers with exception of outliers (beyond 1.5 times interquartile range), as well as the mean. Details of statistical test and *p*-values are reported in the figure legends and in Extended Data Table 1.

---

**Extended Data Figure 1.**
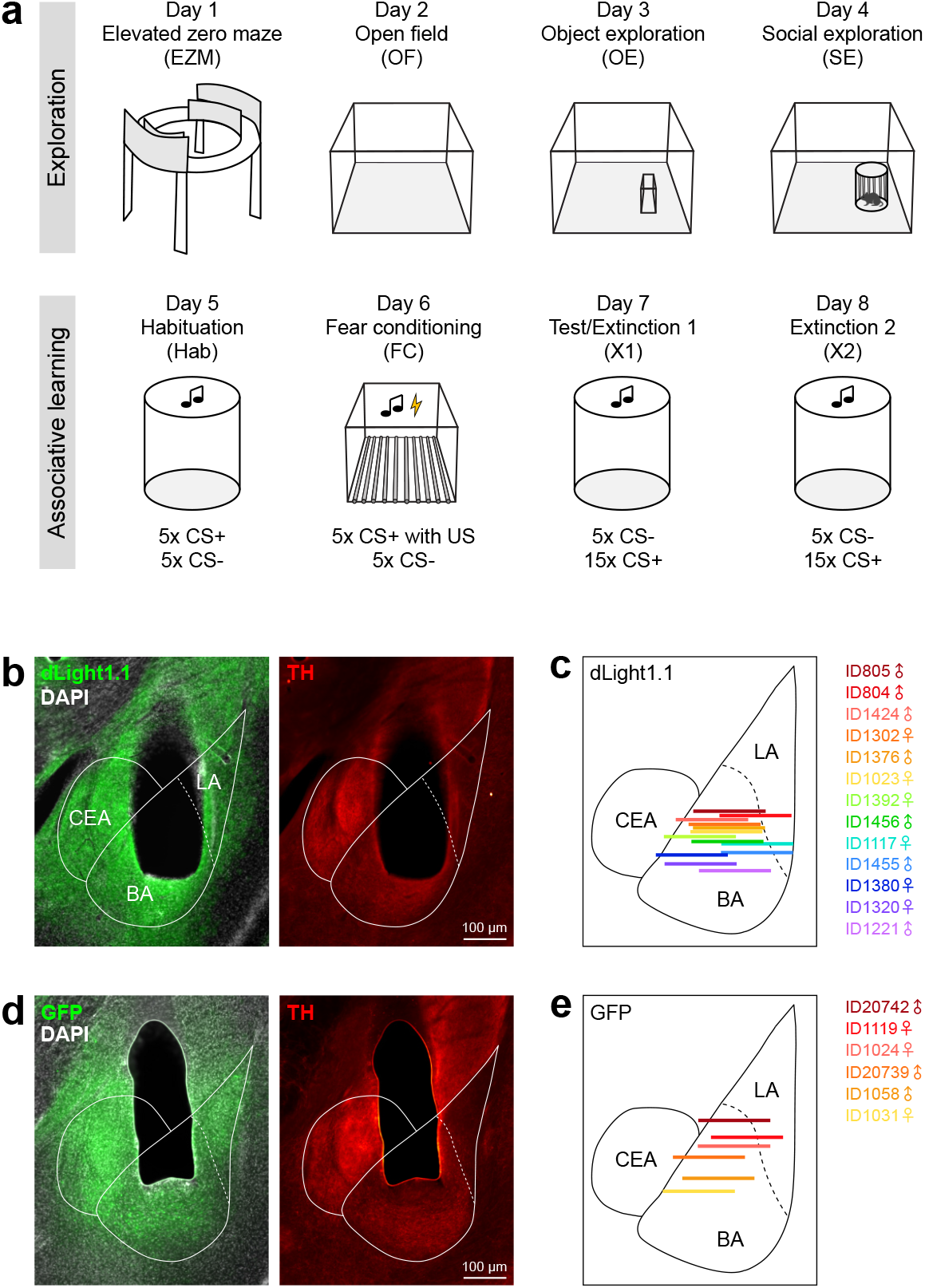
Experimental timeline and fibre placements. **a** Sequence of behavioural paradigms. Tests were performed on consecutive days as detailed in the Methods section. **b** Representative implant site for dLight1.1 recordings (ID1221) showing dLight expression and tyrosine hydroxylase (TH)-positive fibres in the basolateral amygdala (BLA). **c** Schematic illustrating all reconstructed implant sites of fibres within the BLA for dLight1.1 recordings matched to a mouse brain atlas (N=13 mice). Symbols indicate subject sex (male: ♂, female: ♀). **d** Representative implant site for GFP controls (ID1058) showing GFP expression and TH-positive fibres in the BLA. **e** Schematic illustrating all reconstructed implant sites of fibres within the BLA for GFP controls matched to a mouse brain atlas (N=6 mice). LA, lateral amygdala; BA, basal amygdala; CEA, central amygdala; TH, tyrosine hydroxylase.

**Extended Data Figure 2.**
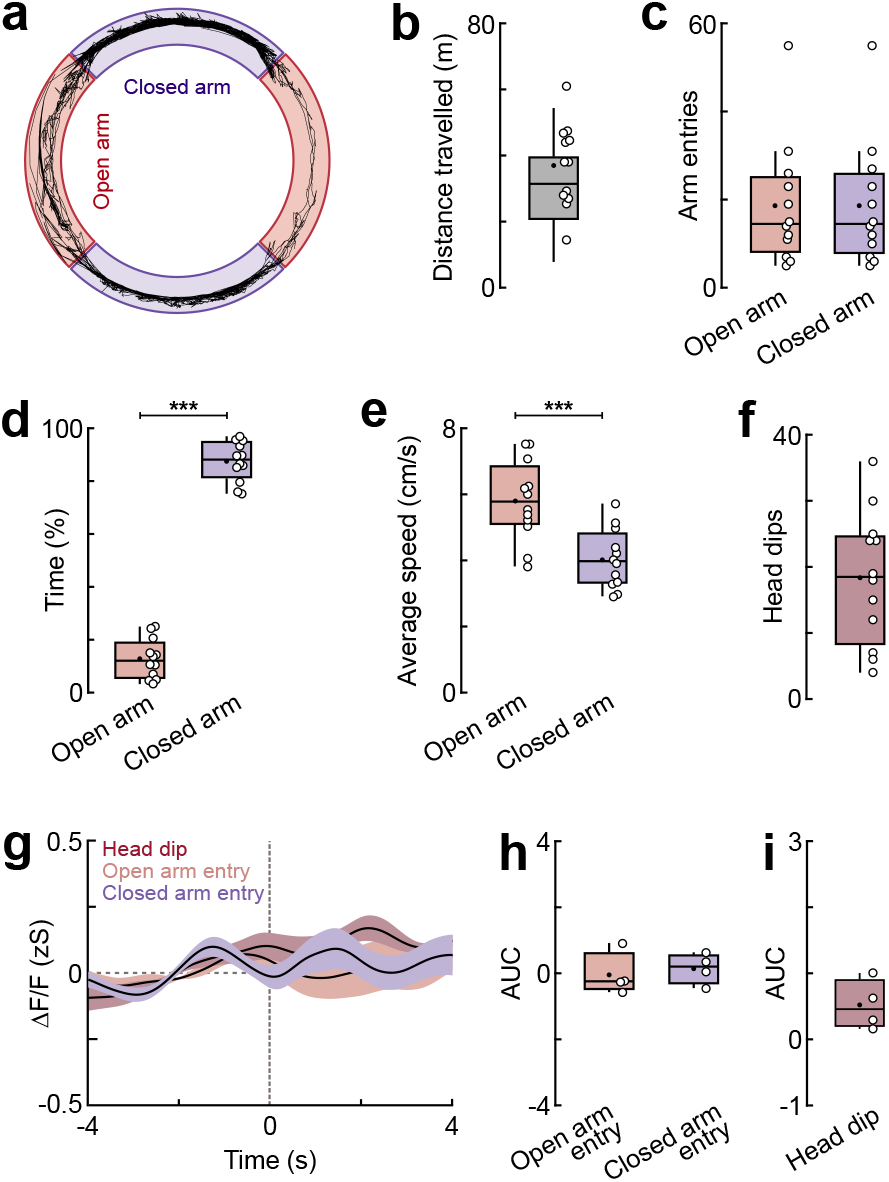
Behavioural measures and control recordings in the elevated zero maze. **a** Track of the example mouse in the elevated zero maze (EZM) shown in Figure 1 for the entire 20-min session. Colour code indicates open (red) and closed arms (purple). Note that within the closed arms, body tracking was less precise because the maze walls partially obscured the animal from the overhead camera. **b** Average distance travelled in the EZM (N=12 mice). **c** Number of open and closed arm entries (N=12). Note that numbers of open and closed arm entries are almost identical for individual mice given the layout of the zero maze. **d** Average time spent in the open and closed arms. Paired t test, *t*(11)=17.28, *p*<0.0001; N=12. **e** Average speed in the open and closed arms. Paired t test, *t*(11)=5.211, *p=*0.0003; N=12. **f** Number of head dip events (N=12). **g** Fluorescence signal in GFP controls time-locked to head dips, open and closed arm entries (N=4). **h** Average area under the curve (AUC) for open vs. closed arm entries in GFP controls (N=4). **i** Average AUC for head dip events in GFP controls (N=4). Average traces in g are weighted means across animals with s.e.m.; Tukey box-and-whisker plots in b-f, h-i show median values, 25^th^ and 75^th^ percentiles, and min to max whiskers with exception of outliers, dots indicate the mean, circles represent individual animals. \*\*\**p*<0.001. Additional details of statistical analyses are provided in Extended Data Table 1.

**Extended Data Figure 3.**
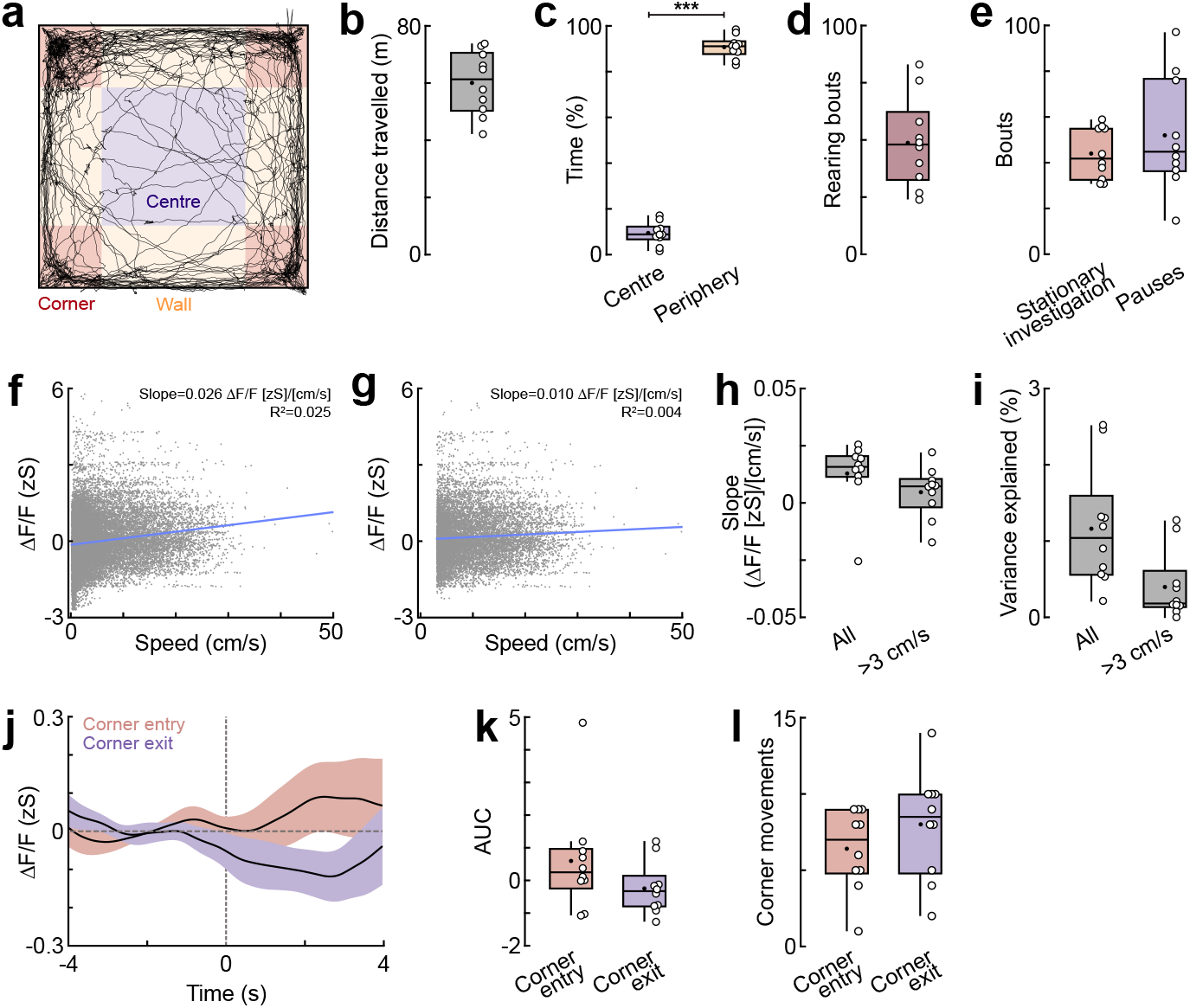
Behavioural measures and dopamine dynamics in the open field. **a** Track of the example mouse in the open field (OF) shown in Figure 1 for the entire 20-min session. Colour code indicates centre (purple), corner (red) and wall areas (yellow). **b** Average distance travelled in the OF (N=10). **c** Average time spent in the centre vs periphery; periphery includes corner and wall areas. Paired t test, *t*(9)=26.45, *p*<0.0001; N=10 mice. **d** Number of rearing bouts (N=10). **e** Number of stationary investigation and pause events (N=10). **f** Representative linear regression of BLA dopamine signals against locomotor speed across all time points. **g** Representative linear regression from the same mouse after excluding periods with locomotor speeds <3 cm/s, thereby restricting the analysis to active locomotion and minimising the contribution of low-translational-movement behavioural states with distinct dopamine dynamics (see Fig. 1l). **h** Slopes of the linear relationship between BLA dopamine signals and locomotor speed for individual mice, calculated across all time points or after exclusion of locomotor speeds <3 cm/s. For ‘All’: one-sample Wilcoxon test, *W*=35.00, *p*=0.0840; for ‘>3 cm/s’: one-sample Wilcoxon test, *W*=23.00, *p*=0.2754; N=10. **i** Percentage of variance in BLA dopamine signals explained by locomotor speed for individual mice, across all time points or after exclusion of locomotor speeds <3 cm/s (N=10). **j** Dopamine release time-locked to corner entries and corner exits (N=10). Corner entries and exits were only scored for transitions between the corner and centre regions, with the centre region defined to include a 4.6-cm margin from the adjacent wall regions for this analysis. Building on previous work showing that the BLA organises activity according to behavioural state rather than spatial position of the animal within the OF, we interpret corner entries as exploratory transitions, typically accompanied by investigation and rearing, whereas corner exits correspond to shifts toward non-exploratory states^4^. **k** Average area under the curve (AUC) for corner transitions. Wilcoxon matched-pairs signed rank test, *W*=-27.00, *p*=0.1934; N=10. **l** Number of corner movements (N=10). Average traces in j are weighted means across animals with s.e.m.; Tukey box-and-whisker plots in b-e, h-i and k-l show median values, 25^th^ and 75^th^ percentiles, and min to max whiskers with exception of outliers, dots indicate the mean, circles represent individual animals. \*\*\**p*<0.001. Additional details of statistical analyses are provided in Extended Data Table 1.

**Extended Data Figure 4.**
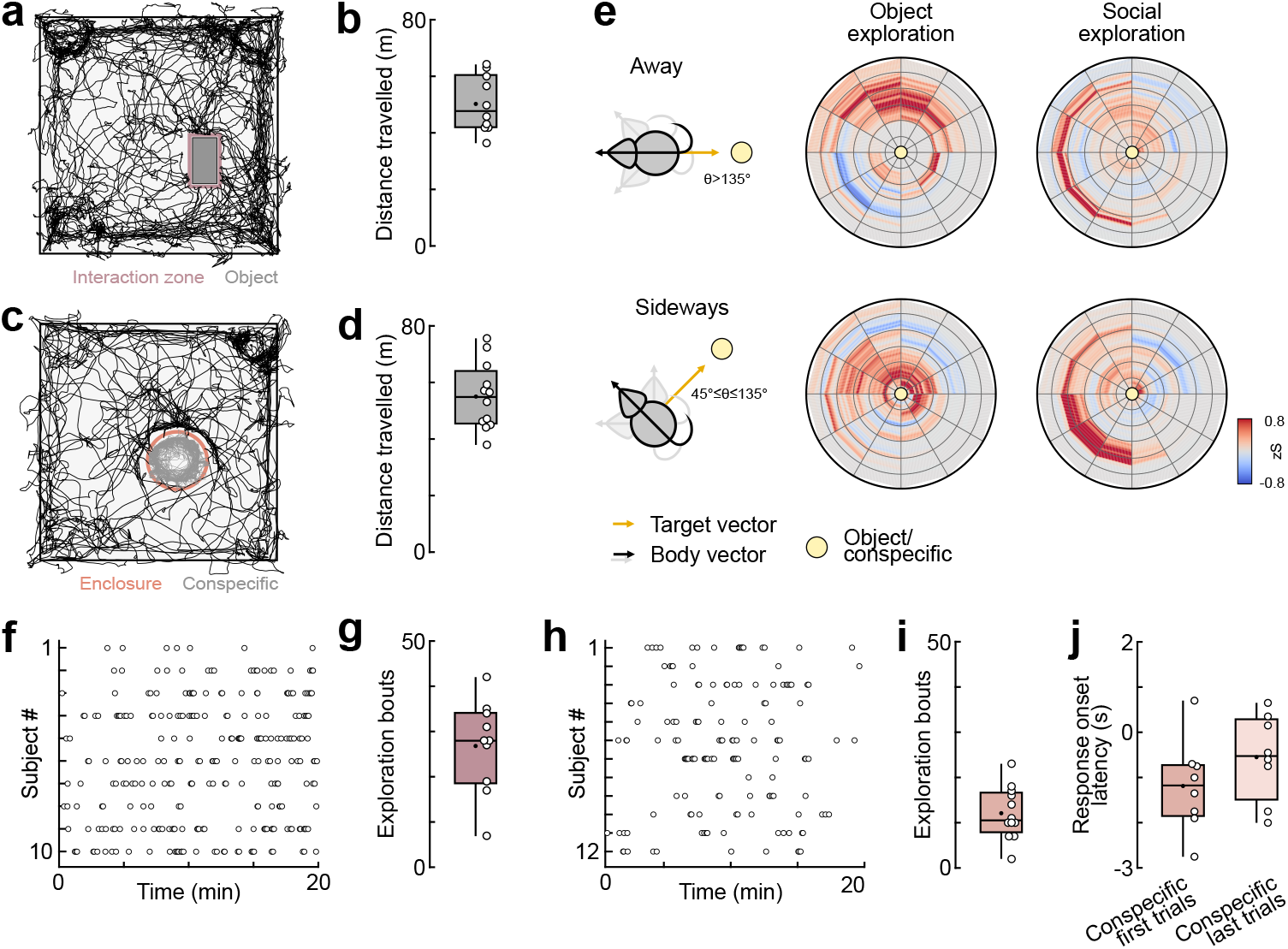
Behavioural measures and dopamine dynamics during stimulus-directed exploration. **a** Track of the example mouse in object exploration (OE) shown in Figure 2 for the entire 20-min session. **b** Average distance travelled in the OE session (N=10 mice). **c** Track of the example mouse in social exploration (SE) shown in Figure 2 for the entire 20-min session. **d** Average distance travelled in the SE session (N=12). **e** Representative polar heatmaps for the “away” and “sideways” conditions, related to Figure 2 (same mouse). The orientation angle θ was computed as the angle between the animal’s body vector (yellow arrow) and the target vector (black vector) pointing to the centre of the object or the enclosure (represented as yellow circle). For the “away” category, only frames with an orientation angle θ>135° are included. For “sideways”, only frames with an orientation angle 45°≤θ≤135° are included. Maps are constructed by converting each frame into polar coordinates relative to the stimulus and binning the z-scored signal by angular sector and radial distance from the target’s centre (radius 30 cm). **f** Raster plot of individual nose-to-object interaction events across mice over time (N=10). **g** Number of nose-to-object events across mice (N=10). **h** Raster plot of individual nose-to-nose interaction events across mice over time (N=12). **i** Number of nose-to-nose events across mice (N=12). **j** Response onset for nose-to-nose contact, defined as the first time point at which the signal reached 10% of its peak amplitude (t_10_), relative to contact onset (0 s). Negative values indicate pre-contact onset during close approach. Onset latency was calculated only for mice exhibiting a positive response peak within the predefined response window (0-4 s) for both timepoints. Paired t test, *t*(7)=1.248, *p*=0.2520; N=8. Tukey box-and-whisker plots in b, d, g, i and j show median values, 25^th^ and 75^th^ percentiles, and min to max whiskers with exception of outliers, dots indicate the mean, circles represent individual animals. Additional details of statistical analyses are provided in Extended Data Table 1.

**Extended Data Figure 5.**
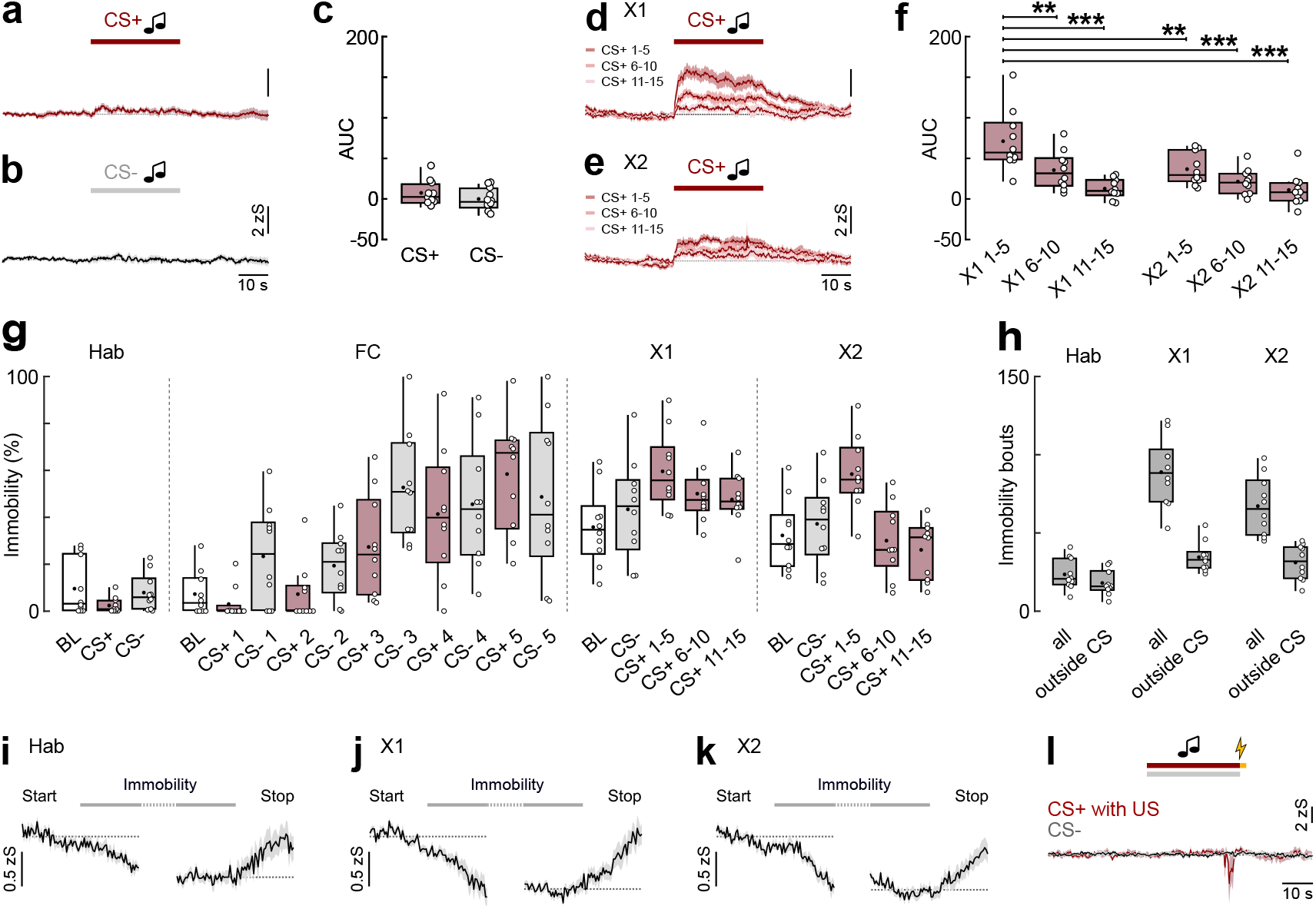
Extended analysis of fear learning and extinction. **a** Basolateral amygdala dopamine release during habituation for CS+ presentations. Shown is the average trace across five trials (N=10 mice). **b** Same for CS−(N=10). **c** Corresponding average area under the curve (AUC) for CS+ and CS−. Paired t test, *t*(9)=1.547, *p*=0.1563; N=10. **d** Development of dopamine release in the Test/X1 session. Average traces are shown as responses to blocks of five CS+ presentations. **e** Same for X2 session (N=10). **f** Corresponding average area under the curve (AUC) for CS+ across X1 and X2 sessions. Repeated-measures one-way ANOVA, *F*(5,45)=11.32, *p<*0.0001, followed by Holm-Šídák multiple comparisons: X1 1-5 vs. X1 6-10, *p*=0.0053; X1 1-5 vs. X1 11-15, *p<*0.0001; X1 1-5 vs. X2 1-5, *p*=0.0081; X1 1-5 vs. X2 6-10, *p<*0.0001; X1 1-5 vs. X2 11-15, *p<*0.0001; all other comparisons n.s.; N=10. **g** Immobility levels throughout the fear conditioning and extinction paradigm (N=10). **h** Number of immobility bouts during habituation, X1 and X2 sessions, separated for the entire session and complete bouts outside any CS presentation used to analyse dopamine dynamics during immobility events (N=10). **i** Average traces of dopamine release during immobility bouts during habituation, aligned to immobility start and stop (N=10). **j** Same for X1 and **k** and X2 sessions. **l** Fluorescence signal for GFP controls during fear conditioning for CS+ and US presentations (red) as well as for CS−control cues (black). Shown is the average trace across five trials (N=6). Average traces in a-b, d-e, and l are arithmetic means across animals with s.e.m.; traces in i-k are weighted means across animals with s.e.m.; Tukey box-and-whisker plots in c, f, g and h show median values, 25^th^ and 75^th^ percentiles, and min to max whiskers with exception of outliers, dots indicate the mean, circles represent individual animals. \*\**p*<0.01, \*\*\**p*<0.001. Additional details of statistical analyses are provided in Extended Data Table 1.

**Extended Data Table 1:** Statistical results. Summary of all statistical analyses for data presented in main and Extended Data figures.

**Main figures**
| Figure | Panel | Sample size | Statistical test | Post hoc comparisons |
| --- | --- | --- | --- | --- |
| 1 | e | N=12 | Paired t test<br>$t(11)=5.716, p<0.0001$ | – |
| 1 | f | N=12 | Linear mixed-effects model, Type III ANOVA with Satterthwaite's approximation<br>Speed bin: $F(3,71)=1.330, p=0.2716$<br>Arm: $F(1,71)=54.02, p<0.0001$<br>Speed bin × arm: $F(3,71)=0.651, p=0.5847$ | – |
| 1 | g | N=12 | One-sample t test<br>$t(11)=4.651, p=0.0007$ | – |
| 1 | j | N=10 | One-sample t test<br>$t(9)=12.40, p<0.0001$ | – |
| 1 | k | N=10 | Paired t test<br>$t(9)=4.486, p=0.0015$ | – |
| 2 | f | N=10 | Paired t test<br>$t(9)=6.051, p=0.0002$ | – |
| 2 | g | N=12 | Paired t test<br>$t(11)=3.149, p=0.0093$ | – |
| 2 | k | N=10 | One-sample Wilcoxon test<br>$W=49.00, p=0.0098$ | – |
| 2 | l | N=12 | One-sample t test<br>$t(11)=1.692, p=0.1188$ | – |
| 2 | o | N=10 object,<br>N=11 conspecific | Unpaired t test<br>$t(19)=3.144, p=0.0053$ | – |
| 2 | p | N=9 | Paired t test<br>$t(8)=2.033, p=0.0765$ | – |
| 3 | b | N=10 | Repeated-measures one-way ANOVA<br>$F(2,18)=31.15, p<0.0001$ | Holm-Šidák multiple comparisons<br>US vs. CS+, $p=0.0722$<br>US vs. CS-, $p<0.0001$<br>CS+ vs. CS-, $p<0.0001$ |
| 3 | d | N=10 | Paired t test<br>$t(9)=3.471, p=0.0070$ | – |
| 3 | f | N=10 | Paired t test<br>$t(9)=4.151, p=0.0025$ | – |
| 3 | h | N=10 | Paired t test<br>$t(9)=2.301, p=0.0470$ | – |
| 3 | k | N=10 | Repeated-measures one-way ANOVA<br>$F(3,27)=19.08, p<0.0001$ | Holm-Šidák multiple comparisons<br>After cond. CS+ vs. After cond. CS-, $p<0.0001$<br>After cond. CS+ vs. After ext. CS+, $p<0.0001$<br>After cond. CS+ vs. After ext. CS-, $p<0.0001$<br>After cond. CS- vs. After ext. CS+, $p=0.7277$<br>After cond. CS- vs. After ext. CS-, $p=0.7277$<br>After ext. CS+ vs. After ext. CS-, $p=0.4683$ |
| 3 | m | N=10 | Paired t test<br>$t(9)=5.425, p=0.0004$ | – |

Extended Data figures
| Figure | Panel | Sample size | Statistical test | Individual comparisons |
| --- | --- | --- | --- | --- |
| ED2 | d | N=12 | Paired t test<br>$t(11)=17.28, p<0.0001$ | – |
| ED2 | e | N=12 | Paired t test<br>$t(11)=5.211, p=0.0003$ | – |
| ED3 | c | N=10 | Paired t test<br>$t(9)=26.45, p<0.0001$ | – |
| ED3 | h | N=10 | All: one-sample Wilcoxon test<br>$W=35.00, p=0.0840$<br>>3 cm/s: one-sample Wilcoxon test<br>$W=23.00, p=0.2754$ | – |
| ED3 | k | N=10 | Wilcoxon matched-pairs signed rank test<br>$W=-27.00, p=0.1934$ | – |
| ED4 | j | N=8 | Paired t test<br>$t(7)=1.248, p=0.2520$ | – |
| ED5 | c | N=10 | Paired t test<br>$t(9)=1.547, p=0.1563$ | – |
| ED5 | f | N=10 | Repeated-measures one-way ANOVA<br>$F(5,45)=11.32, p<0.0001$ | Holm-Šidák multiple comparisons<br>X1 1-5 vs. X1 6-10, $p=0.0053$<br>X1 1-5 vs. X1 11-15, $p<0.0001$<br>X1 1-5 vs. X2 1-5, $p=0.0081$<br>X1 1-5 vs. X2 6-10, $p<0.0001$<br>X1 1-5 vs. X2 11-15, $p<0.0001$<br>X1 6-10 vs. X1 11-15, $p=0.1706$<br>X1 6-10 vs. X2 1-5, $p>0.9999$<br>X1 6-10 vs. X2 6-10, $p=0.6615$<br>X1 6-10 vs. X2 11-15, $p=0.1196$<br>X1 11-15 vs. X2 1-5, $p=0.127$<br>X1 11-15 vs. X2 6-10, $p=0.9408$<br>X1 11-15 vs. X2 11-15, $p>0.9999$<br>X2 1-5 vs. X2 6-10, $p=0.5696$<br>X2 1-5 vs. X2 11-15, $p=0.0871$<br>X2 6-10 vs. X2 11-15, $p=0.8838$ |

## REFERENCES

1. Mehlhorn, K. et al. Unpacking the exploration– exploitation tradeoff: A synthesis of human and animal literatures. Decision 2, 191–215 (2015).

2. Cohen, J. D., McClure, S. M. & Yu, A. J. Should I stay or should I go? How the human brain manages the trade-off between exploitation and exploration. Philos Trans R Soc Lond B Biol Sci 362, 933–942 (2007).

3. Duvarci, S. & Pare, D. Amygdala Microcircuits Controlling Learned Fear. Neuron 82, 966–980 (2014).

4. Gründemann, J. et al. Amygdala ensembles encode behavioral states. Science 364, eaav8736 (2019).

5. Fustiñana, M. S., Eichlisberger, T., Bouwmeester, T., Bitterman, Y. & Lüthi, A. State-dependent encoding of exploratory behaviour in the amygdala. Nature 592, 267–271 (2021).

6. Lutas, A. et al. State-specific gating of salient cues by midbrain dopaminergic input to basal amygdala. Nat Neurosci 22, 1820–1833 (2019).

7. Sias, A. C. et al. Dopamine projections to the basolateral amygdala drive the encoding of identity-specific reward memories. Nat Neurosci 27, 728–736 (2024).

8. Tang, W., Kochubey, O., Kintscher, M. & Schneggenburger, R. A VTA to Basal Amygdala Dopamine Projection Contributes to Signal Salient Somatosensory Events during Fear Learning. J. Neurosci. 40, 3969–3980 (2020).

9. Brickner, M. A., Szot, W. E., Wolff, A. R., Thomas, M. J. & Saunders, B. T. Basolateral amygdala dopamine transmits emotional salience. Nat Commun 17, 7623 (2026).

10. Patriarchi, T. et al. Ultrafast neuronal imaging of dopamine dynamics with designed genetically encoded sensors. Science 360, eaat4422 (2018).

11. Menegas, W., Babayan, B. M., Uchida, N. & Watabe-Uchida, M. Opposite initialization to novel cues in dopamine signaling in ventral and posterior striatum in mice. eLife 6, e21886 (2017).

12. Akiti, K. et al. Striatal dopamine explains novelty-induced behavioral dynamics and individual variability in threat prediction. Neuron 110, 3789–3804.e9 (2022).

13. Salinas-Hernández, X. I., Zafiri, D., Sigurdsson, T. & Duvarci, S. Functional architecture of dopamine neurons driving fear extinction learning. Neuron 111, 3854–3870.e5 (2023).

14. Iordanova, M. D., Yau, J. O.-Y., McDannald, M. A. & Corbit, L. H. Neural substrates of appetitive and aversive prediction error. Neuroscience & Biobehavioral Reviews 123, 337–351 (2021).

15. Grewe, B. F. et al. Neural ensemble dynamics underlying a long-term associative memory. Nature 543, 670–675 (2017).

16. Krabbe, S. et al. Adaptive disinhibitory gating by VIP interneurons permits associative learning. Nat Neurosci 22, 1834–1843 (2019).

17. Lutas, A., Fernando, K., Zhang, S. X., Sambangi, A. & Andermann, M. L. History-dependent dopamine release increases cAMP levels in most basal amygdala glutamatergic neurons to control learning. Cell Reports 38, 110297 (2022).

18. Mocellin, P. et al. A septal-ventral tegmental area circuit drives exploratory behavior. Neuron 112, 1020–1032.e7 (2024).

19. Muller, J. F., Mascagni, F. & McDonald, A. J. Dopaminergic innervation of pyramidal cells in the rat basolateral amygdala. Brain Struct Funct 213, 275–288 (2009).

20. Pinard, C. R., Muller, J. F., Mascagni, F. & McDonald, J. Dopaminergic innervation of interneurons in the rat basolateral amygdala. Neuroscience 157, 850–863 (2008).

21. Chu, H.-Y., Ito, W., Li, J. & Morozov, A. Target-Specific Suppression of GABA Release from Parvalbumin Interneurons in the Basolateral Amygdala by Dopamine. J. Neurosci. 32, 14815–14820 (2012).

22. Aksoy-Aksel, A., Gall, A., Seewald, A., Ferraguti, F. & Ehrlich, I. Midbrain dopaminergic inputs gate amygdala intercalated cell clusters by distinct and cooperative mechanisms in male mice. eLife 10, e63708 (2021).

23. Bulumulla, C. et al. Synapse-like Specializations at Dopamine Release Sites Orchestrate Efficient and Precise Neuromodulatory Signaling. bioRxiv (2026). Preprint at 10.1101/2024.09.16.613338

24. Marowsky, A., Yanagawa, Y., Obata, K. & Vogt, K. E. A Specialized Subclass of Interneurons Mediates Dopaminergic Facilitation of Amygdala Function. Neuron 48, 1025–1037 (2005).

25. Bissière, S., Humeau, Y. & Lüthi, A. Dopamine gates LTP induction in lateral amygdala by suppressing feedforward inhibition. Nat Neurosci 6, 587–592 (2003).

26. Kröner, S., Rosenkranz, J. A., Grace, A. A. & Barrionuevo, G. Dopamine Modulates Excitability of Basolateral Amygdala Neurons In Vitro. Journal of Neurophysiology 93, 1598–1610 (2005).

27. Fink, J. S. & Smith, G. P. Mesolimbic and mesocortical dopaminergic neurons are necessary for normal exploratory behavior in rats. Neuroscience Letters 17, 61–65 (1980).

28. Fink, J. S. & Smith, G. P. Mesolimbicocortical dopamine terminal fields are necessary for normal locomotor and investigatory exploration in rats. Brain Research 199, 359–384 (1980).

29. Morel, C. et al. Midbrain projection to the basolateral amygdala encodes anxiety-like but not depression-like behaviors. Nat Commun 13, 1532 (2022).

30. Nguyen, C. et al. Nicotine inhibits the VTA-to-amygdala dopamine pathway to promote anxiety. Neuron 109, 2604–2615.e9 (2021).

31. Vander Weele, C. M. et al. Dopamine enhances signal-to-noise ratio in cortical-brainstem encoding of aversive stimuli. Nature 563, 397–401 (2018).

32. Paxinos, G. & Franklin, K. Paxinos and Franklin’s the Mouse Brain in Stereotaxic Coordinates. (Academic Press, 2019).

33. Labouesse, M. A., Cola, R. B. & Patriarchi, T. GPCR-Based Dopamine Sensors—A Detailed Guide to Inform Sensor Choice for In Vivo Imaging. International Journal of Molecular Sciences 21, 8048 (2020).

34. Mathis, A. et al. DeepLabCut: markerless pose estimation of user-defined body parts with deep learning. Nat Neurosci 21, 1281–1289 (2018).

35. He, K., Zhang, X., Ren, S. & Sun, J. Deep Residual Learning for Image Recognition. arXiv (2015). Preprint at 10.48550/arXiv.1512.03385

36. Gabriel, C. J. et al. BehaviorDEPOT is a simple, flexible tool for automated behavioral detection based on markerless pose tracking. eLife 11, e74314 (2022).

37. Bruno, C. A. et al. pMAT: An open-source software suite for the analysis of fiber photometry data. Pharmacology Biochemistry and Behavior 201, 173093 (2021).

